# Identification of omega-3 oxylipins in human milk-derived extracellular vesicles with pro-resolutive actions in gastrointestinal inflammation

**DOI:** 10.1101/2023.08.04.551608

**Authors:** Marta Gómez-Ferrer, Elena Amaro-Prellezo, Abel Albiach-Delgado, Isabel Ten-Domenech, Julia Kuligowski, Pilar Sepúlveda

## Abstract

Premature infants (PIs) are at risk of suffering necrotizing enterocolitis (NEC), and infants consuming human milk (HM) show a lower incidence than infants receiving formula. The composition of HM has been studied in depth, but the lipid content of HM-derived small extracellular vesicles (HM sEVs) remains unexplored. We isolated HM sEVs from HM samples and analyzed their oxylipin content using liquid chromatography coupled to mass spectrometry, which revealed the presence of anti-inflammatory oxylipins. We then examined the efficacy of a mixture of these oxylipins in combating inflammation and fibrosis, in vitro and and in a murine model of inflammatory bowel disease (IBD). HM-related sEVs contained higher concentrations of oxylipins derived from docosahexaenoic acid, an omega-3 fatty acid. Three anti-inflammatory oxylipins, 14-HDHA, 17-HDHA, and 19,20-DiHDPA (ω3 OXLP), demonstrated similar efficacy to HM sEVs in preventing cell injury, inducing re-epithelialization, mitigating fibrosis, and modulating immune responses. Both ω3 OXLP and HM sEVs effectively reduced inflammation in IBD-model mice, preventing colon shortening, infiltration of inflammatory cells and tissue fibrosis. Incorporating this unique cocktail of oxylipins into fortified milk formulas might reduce the risk of NEC in PIs and also provide immunological and neurodevelopmental support.

## 1. INTRODUCTION

Human milk (HM) has several nutritional and immunological benefits that favor the clinical evolution and neurodevelopment of premature infants (PIs) in the short- and long term [1]. PIs fed HM, especially their own mother’s milk, are at significantly less risk of serious diseases such as necrotizing enterocolitis (NEC), neonatal sepsis, bronchopulmonary dysplasia and retinopathy of prematurity [2]. However, PIs have higher nutrient requirements than full-term infants, and need enriched milk formulations to meet their nutritional needs, and it is challenging to fulfil their high and variable nutrient requirements during hospitalization [3].

HM consists of 87% water, 1% protein, 4% lipids, and 7% carbohydrates (including 1 to 2.4% oligosaccharides) [4]. It also contains many minerals and vitamins. HM is unique in its high abundance of long-chain polyunsaturated fatty acids (LC-PUFAs), which are derived from two essential fatty acids: linoleic acid (LA, omega-6 [ω6]) and alpha-linolenic acid (ALA, ω3). Elongation of these two LC-PUFAs gives rise to arachidonic acid (AA, ω6) and eicosapentaenoic acid (EPA, ω3), respectively, with the latter further metabolized to docosahexaenoic acid (DHA, ω3) [5]. LC-PUFAs are important for regulating growth, immune function, vision, cognitive development, and motor systems in newborns [6], [7], [8]. There is accumulating evidence that milk-derived bioactive lipids have multifunctional properties [6]. Oxylipins are a diverse class of specialized signaling molecules derived from LC-PUFAs that regulate neonatal intestinal development and protect PIs against intestinal injury [9], [10]. Both ω-3 and ω-6 oxylipins are involved in the initiation and resolution of inflammatory processes [10]. In addition, some oxylipins are precursors of specialized pro-resolving and cytoprotective mediators (SPMs), in particular, ω3-derived oxylipins (resolvins, maresins, and protectins) with anti-inflammatory effects, which are involved in the resolution process following tissue injury [11], [12], [13].

Bioactive compounds of HM can also be transferred from mother to child *via* small extracellular vesicles (sEVs) [8], which are lipid bilayer membrane vesicles (50 to 200 nm) containing myriad signaling molecules including proteins, lipids, microRNAs, mRNAs and other biomolecules protected from degradation [14]. sEVs are present in high concentrations in HM and play an important role in inflammation and immune response of the newborns through intracellular communication [15]. It has been reported that sEVs can escape degradation during digestion, reach gut cells, and be transferred to circulation through lymphatic vessels [16], [17]. Moreover, HM sEVs have been reported to enhance re-epithelization of gut cells and inhibit CD4^+^ T cell activation *in vitro* [18], and restore gut homeostasis in a mouse model of ulcerative colitis [19], [20]. The use of HM sEVs in fortification formulas is, however, controversial due to obvious ethical and logistical reasons [21], and well-defined formulations are needed. Accordingly, a better understanding of the composition of sEVs is essential to identify key molecular players and their mechanism of action. While many characteristics of sEVS are under active investigation, such as surface markers [22], protein cargo [18], [23], and miRNA content [24], little is known about the lipid composition of sEVs from HM [25].

In the present study, we used targeted lipidomic analysis to profile oxylipins in HM sEVs purified from 15 breast milk samples donated by healthy volunteers. We then evaluated and compared the efficacy of a combination of the three most abundant oxylipins in HM sEVs in reducing inflammation in a mouse model of colitis with HM sEVs, which confirmed the potent therapeutic value of 14-HDHA, 17-HDHA, and 19,20-DiHDPA (hereafter referred to as ω3 OXLP). Our findings suggest that the ω3 OXLP formulation could serve as a promising dietary supplement for early and intensive nutrition in PIs to prevent NEC.

## 2. MATERIALS AND METHODS

### 2.1. Ethical Statements

All donors gave their informed consent for inclusion. The study was conducted in accordance with the Declaration of Helsinki, and the protocol was approved by the Ethics Committee of the *Hospital Universitari i Politècnic La Fe*, Valencia, Spain (approval numbers 2021-071-1, 2022-748-1 and 2019-289-1).

Animal procedures were approved by the Ethics Committee of the *Hospital Universitari i Politècnic La Fe* (protocol N° 2021/VSC/PEA/0060) according to guidelines from Directive 2010/63/EU of the European Parliament on the protection of animals used for scientific purposes.

### 2.2. Human Samples

HM samples were obtained from lacting women (28–42 years of age) after informed consent. Fifteen volunteers were enrolled at the Human Milk Bank of the University & Polytechnic Hospital La Fe (Valencia, Spain). Buffy coats of healthy donors were obtained from the Centro de Transfusión de la Comunidad Valenciana (Valencia, Spain) after informed consent.

### 2.3. Cell Culture

Caco-2 epithelial cells (isolated from colon tissue) were maintained in Dulbecco’s modified Eagle’s medium (DMEM)-high glucose (Gibco, Thermo Fisher Scientific, Waltham, MA, USA) supplemented with 10% heat-inactivated fetal bovine serum (FBS, Corning, Glendale, AZ, USA) and 100 U/mL penicillin and 100 μg/mL streptomycin (P/S, Sigma-Aldrich, Saint Louis, MO, USA). Caco-2 cells were stimulated with 60 μg/mL of lipopolysaccharide (LPS) from *Escherichia coli* O111:B4 (Sigma-Aldrich, Darmstadt, Germany) in DMEM-high glucose supplemented with 0.5% FBS and 1% P/S for 24 h in the presence or not of HM sEVs or ω3 OXLP. For differentiation experiments, Caco-2 cells (1×10^5^ cells/cm^2^) were added to 8 μm-pore size Transwell^®^ polycarbonate membranes (Corning^®^ Inc., Corning, NY, USA) in complete medium. Upon reaching a confluent monolayer, Caco-2 cells differentiate spontaneously, and after 21 days they show dense microvilli on the apical side, characteristic of small intestinal enterocytes [26].

Fibroblasts were isolated from human skin biopsies and were cultured in DMEM/F12 (Gibco, Thermo Fisher Scientific) supplemented with 10% FBS and 1% P/S. One day before stimulation, cells were seeded in DMEM/F12 serum-free medium supplemented with 1% P/S. Fibroblasts were then stimulated with LPS (10 ng/mL) for 24 h in the presence or not of HM sEVs or ω3 OXLP in the same medium.

Both fibroblasts and Caco-2 cells were cultured under oxygen/glucose deprivation (OGD) conditions in some experiments, which was induced by culture with DMEM without glucose, glutamine, or phenol red (Thermo Fisher Scientific) in a hypoxia chamber at 1.5% O_2_.

Peripheral blood mononuclear cells (PBMCs) were isolated from buffy coats by density gradient centrifugation with Histopaque (Sigma-Aldrich, Darmstadt, Germany), and were cultured in the Rosewell Park Memorial Institute medium (RPMI, Gibco, Thermo-Fisher Scientific) supplemented with 10% FBS, 1 mM pyruvate, 2 mM glutamine and 1% P/S (all from Sigma-Aldrich). Monocytes were isolated as described [27]. To generate monocyte-derived type 1 or type 2 macrophages (Mϕ1 or Mϕ2, respectively), 5 ng/mL recombinant human granulocyte macrophage-colony stimulating factor (rhGM-CSF, Peprotech) or 20 ng/mL recombinant human macrophage-colony stimulating factor (rhM-CSF, Peprotech) were added to cells in complete RPMI medium. Cytokine stimulation was repeated on day 2 and 4. During the last 16 h of culture, 10 ng/mL of LPS and 20 ng/mL of IFNɣ (R&D Systems, Minneapolis, MN, USA) were added to Mϕ1, whereas 10 ng/mL of LPS and 40 ng/mL of IL4 (PeproTech, London, UK) were added to Mϕ2. Under Mϕ1 conditions, HM sEVs or ω3 OXLP were added on day 0 of the differentiation protocol.

### 2.4. sEV Isolation and Characterization

sEVs were isolated using a serial ultracentrifugation protocol [25]. Briefly, HM was centrifuged three times at 3000×g for 10 min at 4°C to remove milk fat and fat globules. After removing the upper fat layer, the liquid was transferred to a 25-mL polycarbonate bottle and centrifugated twice at 10,000×rpm for 1 h at 4°C. Supernatants were filtered manually through a 0.45-μm filter using a syringe. HM sEVs were then concentrated by three of rounds ultracentrifugation at 30,000 rpm for 2 h at 4°C. Samples were filtered through a 0.22-μm filter to maintain sterility. Protein concentration was determined with the Pierce BCA Protein Assay Kit (Thermo Fisher Scientific) to ensure that equal amounts of protein were used for experiments. sEVs were suspended in RIPA buffer (1% NP40, 0.5% deoxycholate, 0.1% sodium dodecyl sulfate in Tris-buffered saline [TBS], Sigma-Aldrich) for western blotting and in PBS for characterization and functional analysis. Nanoparticle tracking analysis (NTA) and electron microscopy were performed as described [28]. Dynamic light scattering (DLS) was performed to determine the size, distribution, surface charge and stability of sEVs. HM sEVs were placed in a cuvette filled with PBS. The zeta potential magnitude (ζ) and the polydispersity index (PDI) of the samples were measured using a DLS detector (Zetasizer Nano ZS DLS detector, Malvern, UK), which was operated in both continuous and discontinuous modes, employing laser doppler micro-electrophoresis. The instrumental conditions for the DLS system, including temperature, acquisition time, measurement position, and attenuator settings, were optimized for accurate measurements. The specific details of the DLS system setup are described in Table 1, which summarizes the parameters for continuous DLS, discontinuous DLS, and Z-potential measurements. The temperature was maintained at 25°C throughout the experiments, and an equilibration time of 120 s was allowed before each measurement. The acquisition time and attenuator settings were automatically adjusted to seek the optimum conditions for data acquisition. The Smoluchowski model with a correction factor of 1.50 F(ka) was employed for zeta potential calculations, and the voltage was set to auto with a maximum value of 150 V.

**Table 1.** Instrumental conditions of the DLS system.

|  |  |  |
| --- | --- | --- |
| **Continuous DLS** | Acquisition time | 3.0 s |
|  | Measurement position | 4.2 mm |
|  | Attenuator | 11 |
| **Discontinuous DLS** | Equilibration time | 120 s |
|  | Measurement angle | 173° (NIBS default) |
|  | Acquisition time | 10.0 s |
|  | Position | Automatic seek for optimum conditions |
|  | Attenuator | Automatic seek for optimum conditions |
| **Z-Potential** | Model | Smoluchowski (1.50 F(ka)) |
|  | Equilibration time | 120 s |
|  | Acquisition time | Automatic seek for optimum conditions |
|  | Attenuation selection | Automatic seek for optimum conditions |
|  | Voltage | Auto (Max. 150 V) |

**Table 2.**
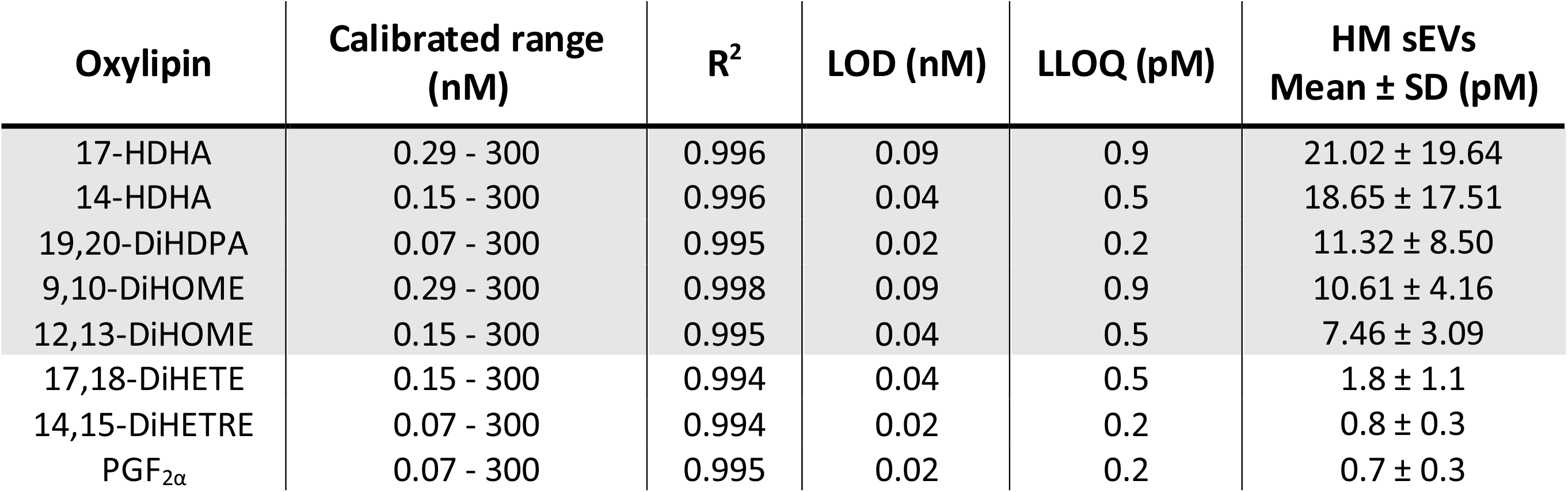
Quantification of oxylipins. Calibration range, linear coefficient of determination (R2), limit of detection (LOD), lower limit of quantification (LLOQ), mean concentration in HM sEVs.

| Oxylipin | Calibrated range (nM) | R<sup>2</sup> | LOD (nM) | LLOQ (pM) | HM sEVs Mean ± SD (pM) |
| --- | --- | --- | --- | --- | --- |
| 17-HDHA | 0.29 - 300 | 0.996 | 0.09 | 0.9 | 21.02 ± 19.64 |
| 14-HDHA | 0.15 - 300 | 0.996 | 0.04 | 0.5 | 18.65 ± 17.51 |
| 19,20-DiHDPA | 0.07 - 300 | 0.995 | 0.02 | 0.2 | 11.32 ± 8.50 |
| 9,10-DiHOME | 0.29 - 300 | 0.998 | 0.09 | 0.9 | 10.61 ± 4.16 |
| 12,13-DiHOME | 0.15 - 300 | 0.995 | 0.04 | 0.5 | 7.46 ± 3.09 |
| 17,18-DiHETE | 0.15 - 300 | 0.994 | 0.04 | 0.5 | 1.8 ± 1.1 |
| 14,15-DiHETRE | 0.07 - 300 | 0.994 | 0.02 | 0.2 | 0.8 ± 0.3 |
| PGF<sub>2α</sub> | 0.07 - 300 | 0.995 | 0.02 | 0.2 | 0.7 ± 0.3 |

### 2.5. Western Blot Analysis

HM sEVs were lysed in 100 μL of RIPA buffer containing protease and phosphatase inhibitors (Complete Mini and PhosSTOP, Sigma-Aldrich). Equal amounts of proteins were mixed with non-reducing Laemmli sample buffer (BioRad) and denatured at 96°C for 5 min. Proteins were separated on 10% SDS-polyacrylamide gels, transferred to polyvinylidene difluoride membranes (Immobilon-P; Millipore), and blocked with TBS containing 5% (w/v) non-fat dry milk powder with 0.1% Tween-20. Human primary antibodies used for western blotting (WB) were as follows: anti-calnexin (dilution 1/1000, Santa Cruz Biotechnology, H-70), anti-Hsp70 (dilution 1/500; Cell Signaling Technology; D69), anti CD63 (dilution 1/500; Santa Cruz Biotechnology; H-193), anti-TSG101 (dilution 1/200; Santa Cruz Biotechnology; C-2), anti-CD81 (dilution 1/500; Santa Cruz Biotechnology; B-11) and anti-CD9 (dilution 1/500; Santa Cruz Biotechnology; C-4). Secondary antibodies were anti-IgG rabbit (dilution 1/4000; Dako; P0448) and anti-IgG mouse (dilution 1/10000; Sigma-Aldrich; A9044). Detection was carried out using peroxidase-conjugated secondary antibodies and the ECL Plus Reagent (GE Healthcare, Chicago, IL, USA) or SuperSignal West Femto (Thermo Fisher Scientific). Reactions were visualized using an Amersham Imager 600 (GE Healthcare) and quantified with ImageJ software (NIH, Bethesda, MD, USA).

### 2.6. Uptake of Labeled HM sEVs

HM sEV uptake by Caco-2 cells was performed after labeling EVs with carboxyfluorescein succinimidyl ester (CFSE; Thermo Fisher Scientific) [29]. Labeled HM sEVs were washed with PBS in an Amicon Ultra-0.5 Centrifugal Filter 100 kDa (Merk, Darmstadt, Germany) and then suspended in 30 µL of filtered PBS and added to 1×10^5^ Caco-2 cells seeded in a 48-well plate. CFSE mixed with PBS was used as a negative control to normalize the amount of unincorporated dye. CFSE-positive cells were detected by flow cytometry after 24 h incubation.

### 2.7. Extraction of the HM-sEVs lipid fraction and oxylipin quantificataion

Sample preparation and oxylipin quantification were adapted as described elsewhere [30]. In short, HM sEVs were extracted using a solid phase extraction Oasis^®^ MAX 96 well plate from Waters (Taunton, MA, USA). Recovered sample extracts were evaporated using a miVac centrifugal vacuum concentrator (Genevac Ltd., Ipswich, UK) and then dissolved in 60 µL methanol:acetonitrile (50:50, v/v).

Sample extracts were analyzed using an Acquity-Xevo TQ-XS system (Waters, Milford, MA, USA) operating in negative electrospray ionization mode. Separations were performed on a Waters Acquity UPLC BEH C18 (2.1×100 mm, 1.7 µm) column using a 0.1% v/v acetic acid and acetonitrile: isopropanol (90:10 v/v) binary gradient. Mass spectrometry (MS) detection was carried out by multiple reaction monitoring. Oxylipins quantified were as follows: 12,13-DiHOME, 9,10-DiHOME, 14,15-DiHETRE, PGE2, PGF2α, 19,20-DiHDPA, 17-HDHA, 14-HDHA, 17,18-DiHETE, 14,15-DiHETE, Resolvin D5, Maresin 2, and 8(S),15(S)-DiHETE.

### 2.8. T-cell Proliferation Assays

T-cell proliferation assays were performed as described [28]. Prior to culturing, PBMCs were labeled with 5 μM CFSE and activated with Dynabeads™ Human T-Activator CD3/CD28 (Thermo Fisher Scientific). To evaluate immunosuppressive potential, we added 15 µg of HM sEVs to 1×10^5^ CFSE-labeled PBMCs in 0.5 mL of medium. After 5 days of culture, T-cell proliferation was determined by flow cytometry to measure CFSE dilution. Analyses of flow cytometry data and the expansion index (EI) [31], were performed with FlowJo software (FlowJo LLC, BD, Franklin Lakes, NJ, USA). The percentage of immunosuppression was calculated by normalizing data to a 0–100% scale by establishing 0% immunosuppression for the EI of activated PBMCs untreated (Act) and 100% immunosuppression for the EI of non-activated PBMCs (No act) using the following formula:

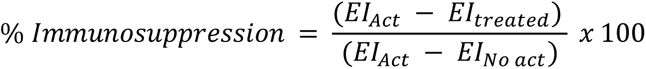

### 2.9. Flow Cytometry

For flow cytometry analysis PBMCs or macrophages were first incubated with a blocking solution (PBS containing 1% normal mouse serum) for 10 min and then incubated with saturating amounts of fluorochrome-conjugated antibodies for 1 h at 4°C. Human antibodies used were as follows: anti-CD3 (PerCP-Cy, BD Biosciences; SK7), anti-CD4 (BV510, BD Biosciences; L200), anti-CD8 (PE-Cy7, BD Biosciences; RPA-T8), anti-CD14 (RPE, Dako, TUK4, Santa Clara, CA, USA), anti-CD163 (PerCP-Cy, BD Biosciences, GHI/61), anti-CD80 (APC, BD Biosciences, FUN-1), anti-CD86 (V450, BD Biosciences, L307.4) and anti-HLA-DR (FITC, Miltenyi Biotec, AC122) at concentrations recommended by the manufacturers. Cells were analyzed on a BD FACSCANTO II flow cytometer equipped with Flowjo^®^ software.

### 2.10. Cell viability assay

To test whether HM sEVs or ω3 OXLP affected cell viability, Caco-2 cells were cultured at a density of 1×10^4^ cells/cm^2^ on a 96-well plate and were then stimulated with LPS or cultured under OGD conditions and treated with HM sEVs or ω3 OXLP. After 24 h the Cell Counting Kit-8 (CCK-8) assay was used to measure proliferation. The optical density of the cultures was measured at 450 nm in each well 4 h after incubation with the CCK-8 assay solution.

### 2.11. Lactate Dehydrogenase Assay

Caco-2 cells were seeded at 1×10^4^ cells/cm^2^ in complete medium. On the next day, cells were cultured under OGD conditions for 24 h, and the supernatant was tested for lactate dehydrogenase using the Cytotoxicity Detection KitPLUS (LDH) (Roche, Indianapolis, IN, USA).

### 2.12. Oxidative Stress Assay

LPS- and OGD-treated cells were washed with PBS and stained with 5 μM 2′,7′-dichlorofluorescin diacetate (DCFH-DA; Sigma-Aldrich) for 20 min at 37°C to detect cell reactive oxygen species (ROS). After staining, cells were washed three times with PBS to eliminate the unconjugated stain, and were detached with trypsin/ethylenediaminetetraacetic acid (EDTA) for flow cytometry.

### 2.13. Scratch Assay

Caco-2 cells and fibroblasts were seeded in a 24-well plate at 2×10^5^ cells/well. Caco-2 cells were stimulated with LPS and treated with HM sEVs or ω3 OXLP for 48 h. To develop scratch assays under OGD conditions, the medium was replaced after 24 h with complete medium and cells were cultured under standard oxygen conditions. Caco-2 cells were then stimulated with LPS and treated with HM sEVs or ω3 OXLP for 48 h. A straight line in the monolayer of cells was created using a 20-μL pipette tip. Images were taken up to 48 h after the addition of treatments using a Leica DM600 inverted microscope at 10× magnification. The area of the scratch wound was then measured using ImageJ.

### 2.14. Real Time Quantitative PCR

RNA was extracted using RLT buffer (Qiagen, Dusseldorf, Germany) and purified with the RNeasy Plus Mini Kit (Qiagen). RNA was quantified spectrophotometrically using a NanoDrop ND-1000 (NanoDrop Technologies, Wilmington, DE, USA). cDNA was obtained by reverse transcription using the PrimeScript RT Reagent Kit (Takara, Kusatsu, Japan). RT-qPCR was performed with the respective human-specific sense and antisense primers and RT-SYBR™ Green PCR Master Mix (Applied Biosystems). Multiwell plates of 384 wells were run on a Viia 7 PCR System (Applied Biosystems). The primers used were:

*hGAPDH* CCCCTCTGCTGATGCCCCA (F) and TGACCTTGGCCAGGGGTGCT (R)

*hTNF-α* CCCTCTGGCCCAGGCAGTCA (F) and ATGGGTGGAGGGGCAGCCTT (R)

*hCOX2* GAATCATTCACCAGGCAAA (F) and TCTGTACTGCGGGTGGAACA (R)

*hOCLN* GGACTGGATCAGGGAATATC (F) and ATTCTTTATCCAAACGGGAG (R)

*hCLDN* CCGGGTTGCCCACCTGCAAA (F) and CGTACATGGCCTGGGCGGTC (R)

*hTGF-β* GAGTGTGGAGACCATCAAGGA (F) and CTGTTTTAGCTGCTGGCGAC (R)

*hIL-1β* AGGCACAAGGCACAACAGGCT (F) and AACAACTGACGCGGCCTGCC (R)

*hIL6* CATTCTGCCCTCGAGCCCACC (F) and GGCAGCAGGCAACACCAGGA (R)

*hIL8* CGTGGCTCTCTTGGCAGCCTTC (F) and TTCCTTGGGGTCCAGACAGAGCTC (R)

*hTLR4* CCCTGCGTGGAGGTGGTTCCTA (F) and CTCCCAGGGCTAAACTCTGGATGGG (R)

*hMMP1* GTGTCTCACAGCTTCCCAGCGAC (F) and GCACTCCACATCTGGGCTGCTTC (R)

*mActβ* GCCAACCGTGAAAAGATGACC (F) and GAGGCATACAGGGACAGCAC (R)

*mArg1* GTGGGGAAAGCCAATGAAGAG (F) and TCAGGAGAAAGGACACAGGTTG (R)

*mCd206* TGTGGAGCAGATGGAAGGTC (F) and TGTCGTAGTCAGTGGTGGTTC (R)

*mCcr2* GTAGTCACTTGGGTGGTGGC (F) and TACAGCGAAACAGGGTGTGG (R)

*mCx3cr1* ACTCCGGTCTCATTTGCAGG (F) and GGGACCTCTGTAGGAGCAGA (R)

*mTnf-α* CCCTCACACTCAGATCATCTTCT (F) and GCTACGACGTGGGCTACAG (R)

*mIl-4* GTACCAGGAGCCATATCCACG (F) and CGTTGCTGTGAGGACGTTTG (R)

*mIl-10* GGACAACATACTGCTAACCGAC (F) and CCTGGGGCATCACTTCTACC (R)

### 2.15. Immunofluorescence analysis

Caco-2 cells were cultured on Transwells^®^ for differentiation. After 21 days, cells were cultured under LPS or OGD conditions and treated with HM sEVs or ω3 OXLP for 24 h. On the next day, cells were fixed in 4% paraformaldehyde for 10 min, washed three times with PBS and permeabilized and blocked with 5% BSA and 0.1% Triton X-100 in PBS for 1 h. Cells were then incubated with mouse anti-human occludin (Santa Cruz, E-5) and rat anti-human E-cadherin (EMD Millipore, DECMA-1) at a concentration of 1/200 in a humidified incubator overnight. Goat anti-mouse IgG (1:500, Alexa Fluor^®^ 488, Abcam) and goat anti-rat IgG (1:500, Alexa Fluor^®^ 555, Abcam) were used as secondary antibodies. Nuclei were stained with DAPI. Quantification of mean fluorescence intensity (MFI) was performed using ImageJ.

### 2.16. Pyrogen test assay

An *in vitro* pyrogen test using PBMCs was used to detect substances that activate human immune cells to express pro-inflammatory cytokines such as TNFα, IL-1β, IL-6 and IL-8 by qPCR. PBMCs (4×10^6^ cells/mL) were incubated with HM sEVs and ω3 OXLP for 5 h. LPS at concentration of 1 µg/mL was used as a positive control.

### 2.17. Mice

Adult male Balb/c mice (6 weeks old, 18−22 g) were purchased from Envigo (Inotiv Inc., Indianapolis, Indiana, USA), and maintained under standard laboratory conditions. All animal procedures were approved by institutional ethical and animal care committees.

### 2.18. TNBS-induced Colitis

2,4,6-trinitrobenzenesulfonic acid (TNBS) was used to induce colitis, a type of IBD, by an intrarectal administration of 100 µL of TNBS (3.5 mg per 20 g mice, Sigma-Aldrich) dissolved in 40% ethanol, as described [32]. The sham group received 100 µL of 0.9% NaCl dissolved in 40% ethanol. Mice were treated by oral gavage with 50 μg of HM sEVs or 0.5 μg of ω3 OXLP prepared in 100 µL of PBS. The untreated TNBS group only received 100 µL of PBS. Treatment was administrated just after colitis induction and at day 1 and 2 thereafter. Mice were sacrificed by cervical dislocation on day 4 after colitis induction. Colon length was measured, and colon tissue was frozen in liquid nitrogen for RNA and protein extraction and fixed in 4% paraformaldehyde acid and embedded in paraffin for immunohistochemistry.

### 2.19. Production of Oxylipins Preparation for *In Vivo* Assays

Oxilipins preparation for *in vivo* assays were prepared as described before [33]. PBS was supplemented with 10% fatty acid-free bovine serum albumin (FAF-BSA, Sigma-Aldrich) and stirred for 3 h at room temperature and then filtered through a 0.22-µm filter. For every dose, 0.5 μg of a mixture of 14 HDHA, 17 HDHA and 19-20 DiHDPA at the same concentration each, was prepared together on 100 μL of PBS and stirred again for 16 h at 37°C. Oxylipins were freshly prepared before experiments. Mice received three doses of ω3 OXLP, a cumulative dose of 1.5 μg/mouse.

### 2.20. Myeloperoxidase activity

Protein was extracted by homogenizing colon tissue, and myeloperoxidase (MPO) activity was determined using a Colorimetric Activity Assay Kit (Sigma-Aldrich, St. Louis, MO, USA), and the optical density of the reaction was measured at 412 nm in a micro-plate reader. MPO activity was expressed as U/μg protein.

### 2.21. Cytokine protein array

Colon samples were homogenized in PBS with protein inhibitors. Samples from each group were pooled and then a BCA assay was performed. A normalized protein content was analyzed with the Proteome Profiler Mouse Cytokine Array Kit, Panel A, (R&D systems, Inc., Minneapolis, Minnesota, USA). The array membrane was blocked for 1 h and then washed. Colon samples and the array detection antibody cocktail were mixed and added to the blocked membrane followed by overnight shaking at 4°C. After washing, streptavidin-HRP buffer was added to the membrane and incubated for 30 min. Following washing, a chemiluminescence reagent mix was added and measurements were performed using an Amersham Imager 600 (GE Healthcare) and quantified with ImageJ.

### 2.22. Measurement of Cytokines by Enzyme-Linked Immunosorbent Assay (ELISA)

Supernatants of the *in vitro* macrophage differentiation, supernatants of colonic tissue homogenized, and the mice plasma were collected and used to measure the levels of TNF-α and IL-10. The quantification of these pro-inflammatory cytokines was conducted by commercial ELISA kits (Invitrogen, Waltham, MA, USA), according to the manufacturer’s instructions.

### 2.23. Mouse Histology and Immunofluorescence

Paraffin-embedded colon samples were cut into 5-µm-thick sections and stained with hematoxylin-eosin (Sigma-Aldrich) to evaluate inflammatory infiltrates, the presence of ulceration and the lesion of crypts. In addition, a blind pathological examination was carried out and tissues were scored using the histological colitis scoring method. This score tests for three tissue characteristics: inflammation severity (grade 0 = none, grade 1 = mild, grade 2 = moderate, and grade 3 = severe), crypt damage (grade 0 = none, grade 1 = basal 1/3 damaged, grade 2 = basal 2/3 damaged, grade 3 = crypts lost and surface epithelium present, and grade 4 = crypts and epithelium lost) and colon wall thickness (grade 0 = normal, grade 1 = medium, grade 2 = large, and grade 3 = very large); all three relativized to the percentage involvement (grade 0 = 0%, grade 1 = 1–25%, grade 2 = 26–50%, grade 3 = 51–75%, and grade 4 = 76–100%). The score pathology was calculated as the sum of each characteristic multiplied by the percent involvement. The total maximum score is 40. A Picro-Sirius Red stain (Direct Red 80 and Picric Acid, Sigma-Aldrich) was used to visualize fibrosis. Slides were visualized on a Leica DMD108 Digital Microscope (Leica Microsystems). For immunofluorescence, slides were blocked with 5% normal goat serum and 0.1% Triton X-100 in PBS for 1 h. Slides were then incubated with rabbit anti-MUC2 (dilution 1/200, Invitrogen, PA5-21329), rat anti-F4/F80 (dilution 1/200, Abcam, ab6640), rabbit anti-CD206 (dilution 1/200; Abcam, ab64693) or rabbit anti-CD274 (dilution 1/200, AB Clonal A11273) overnight in a humidified chamber at 4°C and then slides were washed three times for 5 min each in PBS. Slides were incubated with anti-rat IgG Alexa 555 or anti-rabbit IgG Alexa 488 secondary antibodies for 1 h and then washed three times for 5 min in PBS. Cell nuclei were stained with DAPI (4 ’,6-diamidino-2-fenilindol) and slides were mounted using FluorSave™ Reagent (Merck Millipore). The sections were observed and visualized on a Leica DM2500 fluorescent microscope (Leica Microsystems). Final image processing and quantification were performed with ImageJ by counting green and red spots in the fixed area.

### 2.24. Statistical Analysis

Data are expressed as mean ± SD (standard deviation) or standard error of the mean (SEM), as specified. Student’s t test was used for unpaired samples in between-group comparisons. One-way analysis of variance (ANOVA) was used to compare means of more than 2 groups. A two-way ANOVA was used to simultaneously evaluate the effect of two factors on a response variable. Analyses were conducted with GraphPad Prism 8 software (San Diego, CA, USA). Differences were considered statistically significant at p < 0.05 with a 95% confidence interval.

### 2.25. Data availability

We have submitted all relevant data of our experiments to the EV-TRACK knowledgebase (EV-TRACK ID: EV230969) [34].

## 3. RESULTS

### 3.1. Isolation and Characterization of HM sEVs

sEVs were isolated from HM by sequential centrifugation and filtration [25]. Purified sEVs showed a median number of particles of 1.3 x 10^11^ and a median size of 158 nm, as determined by NTA (Figure 1A). We also used DLS to measure the size of vesicles, the ζ potential (which gives an indication of the potential stability of the colloidal system), and the PDI, which is used to characterize the size distribution of sEVs. The ζ potential was -7.7 ± 1.0 mV, which represents an incipient instability of the system, so it cannot be stored for a long time, or the particles will tend to aggregate. The PDI was 0.390 (Figure 1B), indicating a relatively even size distribution of sEVs. WB revealed that the sEVs expressed the typical markers Hsp70, CD63, TSG101, CD81 and CD9 (Figure 1C), but were negative for the endoplasmic reticulum protein calnexin. Finally, transmission electron microscopy analysis of sEVs revealed a round or cup-shaped morphology and the size was consistent with the findings of NTA (Figure 1D).

**Fig. 1.**
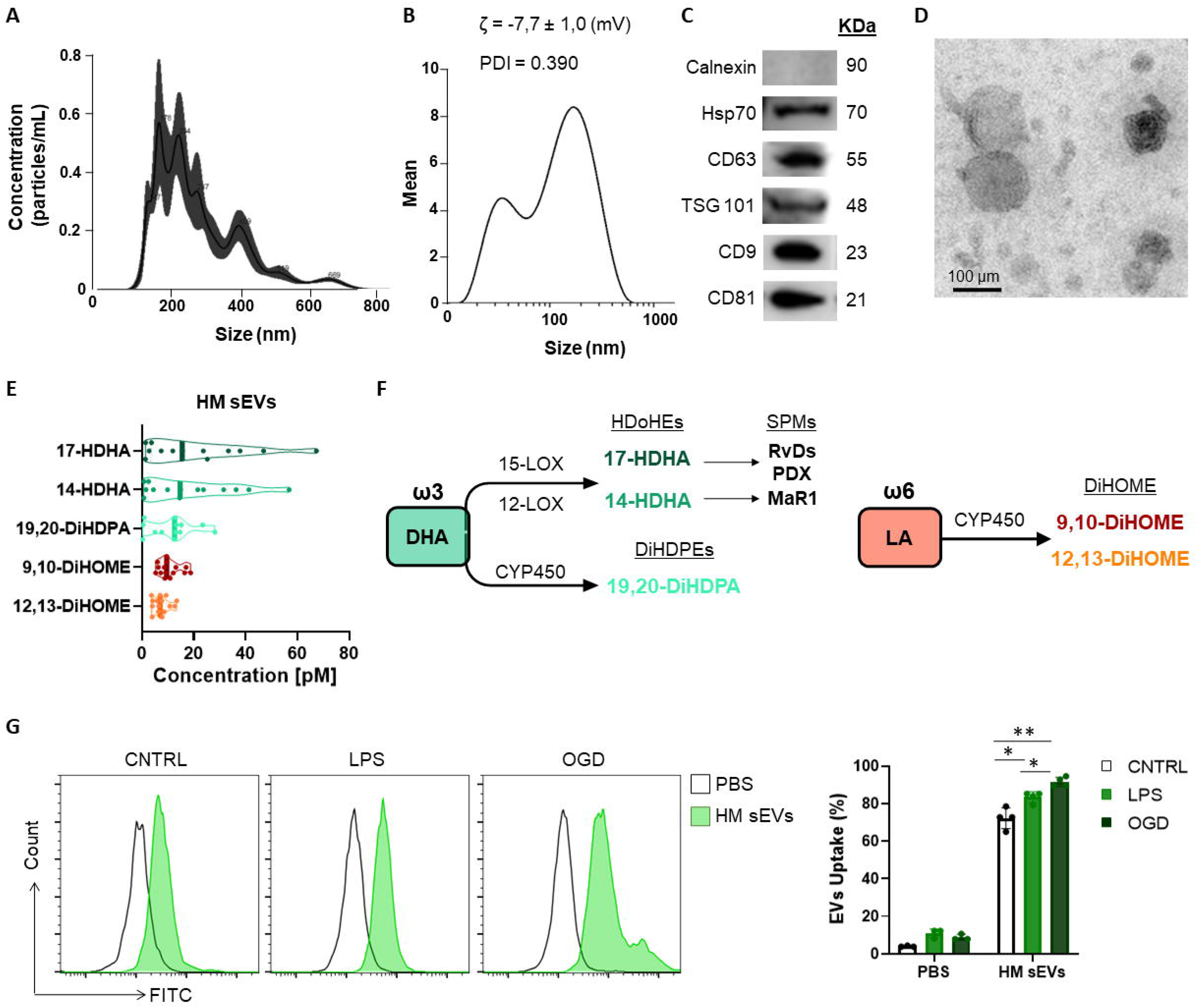
Characterization of HM sEVs and oxylipin content. (A) Representative images of HM sEVs assessed by nanoparticle tracking and (B) DLS analysis; (C) representative western blots of Hsp70, CD63, TSG101, CD81 and CD9 proteins in 30 μg of HM sEVs; absence of calnexin signifies a pure sEVs preparation (D) representative transmission electron microscopy images of HM sEVs. Scale bar: 200 nm; (E) concentration [nM] of the more abundant oxylipins in HM sEVs isolated from 25 mL of HM; (F) scheme of oxylipin synthesis; (G) Intestinal epithelial cells were incubated with CFSE-labeled HM sEVs for 3 h at 37°C and sEV internalization was assessed by flow cytometry. As a negative control, PBS was mixed with CFSE and added to cells in parallel. Representative histograms are shown. sEV internalization was measured by fluorescence intensity and is represented as the percentage of sEV uptake. Graphs represent mean ± SD of four independent experiments. Two-way ANOVA was used for statistical analysis. * p < 0.05, ** p < 0.01.

### 3.2. Quantification and Comparison of Oxylipins in HM sEVs

Quantification of oxylipins from HM sEV samples was performed by means of a validated LC-MS and multiple reaction monitoring. Of the different oxylipins identified, the following could be quantified both in HM sEVs: 9,10-DiHOME, 12,13-DiHOME, 14-HDHA, 17-HDHA, 19,20-DiHDPA. Moreover, 14,15-DiHETE, 14,15-DiHETrE, 17,18-DiHETE, PGE_2_, and PGF_2α_ (Table 1). The most abundant oxylipins were 9,10-DiHOME, 12,13-DiHOME, 19,20-DiHDPA, 14-HDHA and 17-HDHA (Figure 1E). In general, HM-derived sEVs showed higher concentration of DHA-derived oxylipins than LA-derived oxylipins. The former have been reported to have anti-inflammatory activity, and the latter show pro-inflammatory activity (Figure 1E and F) [10]. Based on these results, we investigated whether the protective effects of HM-derived sEVs could be partly attributed to the presence of the three ω3-derived oxylipins – 19,20-DiHDPA, 14-HDHA and 17-HDHA – hereafter referred to as ω3 OXLP.

### 3.3. Protective Effects of HM sEVs and ω3 OXLP on Intestinal Epithelial Cells under Stress and Ischemic Conditions

The main risks for developing NEC are known to be a weak immune system, which increases the presence of infection, and lack of blood flow reaching the colon to supply intestinal cells with oxygen and nutrients, preventing their maturation [35]. To emulate these conditions *in vitro*, Caco-2 intestinal epithelial cells were stimulated with LPS at 60 µg/mL or were cultured in OGD to mimic an ischemic environment. We used an internalization assay with CFSE-stained HM sEVs to question how stress and ischemic conditions affected the uptake of HM sEVs by intestinal cells. Uptake of HM sEVs was observed in 72.3 ± 5.5% of intestinal cells 3 h after their addition to cultures (Figure 1G), and this was increased by 11.2% and 19.3%, respectively, when cells were treated with LPS and OGD (Figure 1G). Notably, cell death increased in Caco-2 cells treated with LPS or OGD, likely due to an increase in cytotoxicity and oxidative stress (ROS) (Figure 2). To assess the protective effect of HM sEVs and ω3 OXLP, Caco-2 cells were treated with 7.5 µg/mL of HM EVs or 0.5 nM of each of the three oxylipins. Both HM sEVs and ω3-OXLP protected Caco-2 cells from LPS-induced damage, improving cell viability over non-treated cells (Figure 2A). Treatment with HM sEVs and ω3 OXLP also decreased cytotoxicity (Figure 2B) and oxidative stress (Figure 2C). Similar results were found under OGD conditions with respect to cell death (Figure 2D). However, only ω3 OXLP treatment had a significant protective effect against cytotoxicity (Figure 2E) and oxidative stress (Figure 2F) generated by OGD. We next tested whether the cell injury triggered by LPS and OGD also affects migration. Indeed, a major concern of PIs with NEC is the presence of "wounds" in the intestine due to the lack of tissue maturation. If the wounds are not repaired the prognosis for the PIs is poor [36]. To investigate whether HM sEVs and ω3 OXLP modulate the migration of intestinal epithelial cells, we used an *in vitro* scratch-wound assay. Results showed that wound closure was slower in cells treated with LPS and OGD, and treatment with HM sEVs or ω3 OXLP restored their migratory capacity and proliferation rate, promoting the development of a continuous monolayer (Figures 2G and 2H).

**Fig. 2.**
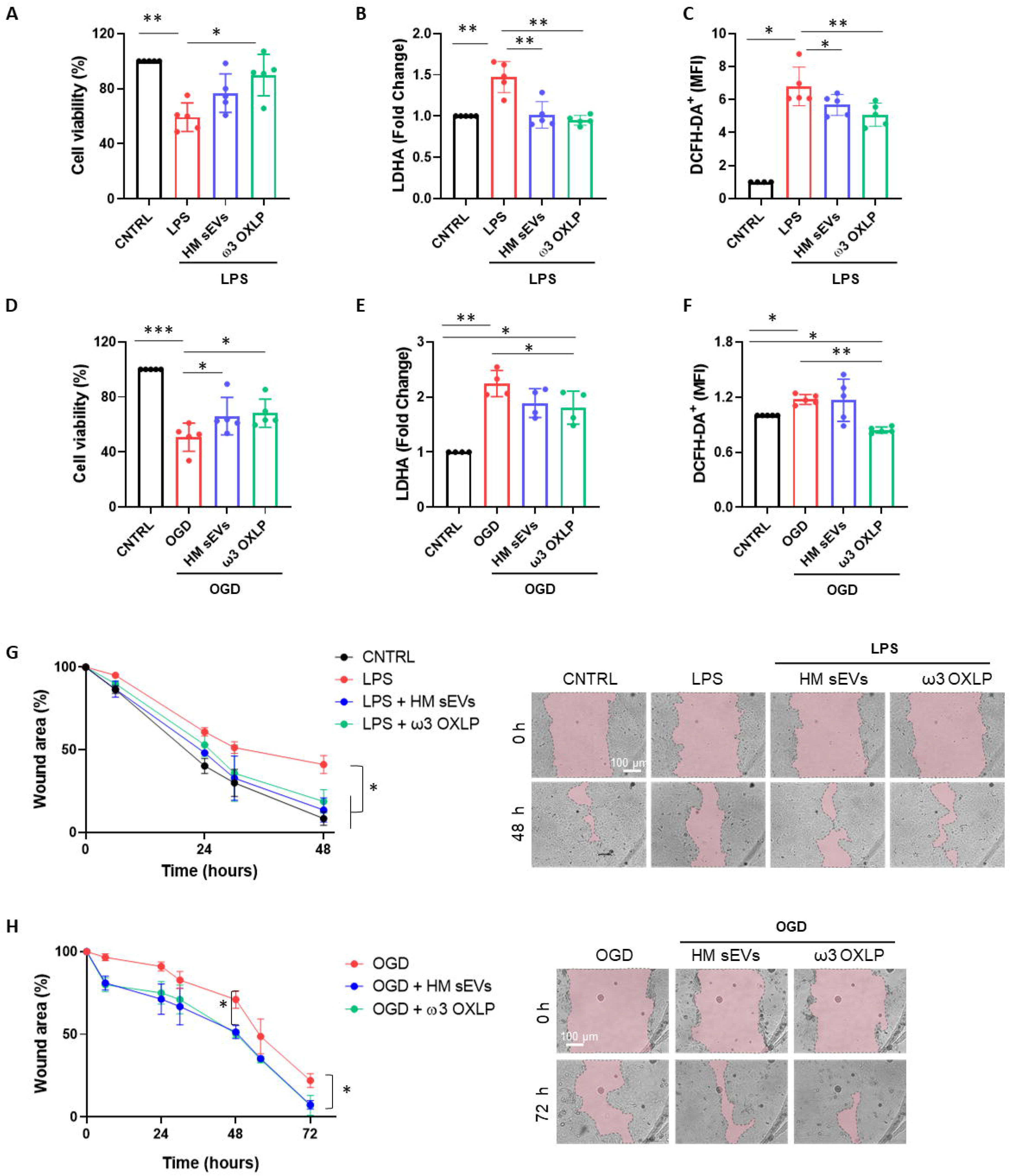
HM-derived sEVs and ω3 oxylipins protect intestinal epithelial cells from damage. (A) Quantification of cell viability measured by CCK8 assay (B); cell cytotoxicity measured by LDH assay and (C) Reactive oxygen species (ROS) production measured by DCFH-DA oxidation in intestinal cells stimulated with lipopolysaccharides (LPS) (100 ng/mL) or oxygen/glucose deprivation (OGD) (D), (E), (F). One-way ANOVA was used for statistical analysis. Quantification of intestinal cell wound area (G) after LPS (100 ng/mL) or OGD (H) treatment. Data were normalized to initial wound area and represented as mean percentage ± SD. One-way ANOVA was used for statistical analysis at different points. Representative brightfield images of wound healing assay at different times (0 and 48 or 72 h) are shown (pink area represents opened wound). Images were taken at 10× magnification. Scale bar: 100 μm. Experiments were performed in triplicate. * p < 0.05, ** p < 0.01.

### 3.4. Modulation of Pro-inflammatory Genes and Tight Junction Proteins by HM sEVs and ω3 OXLP in Inflammatory Conditions

Inflammatory responses triggered by LPS or hypoxia in the intestinal epithelium trigger the upregulation of pro-inflammatory genes such as tumor necrosis factor alpha (*TNF-α*) and cyclooxygenase-2 (*COX-2*). TNF-α plays a critical role in the pathogenesis of inflammatory bowel disease (IBD) by increasing intestinal cell death and detachment in the gut, which damages the integrity of the epithelial barrier [37]. COX-2, an enzyme that accelerates inflammation, also plays a role in the pathophysiological processes of intestinal inflammation [38]. As expected, both LPS and OGD increased the expression of these genes in Caco-2 cells, whereas co-treatment with 7.5 µg/mL of HM sEVs or 0.5 nM of each of the three ω3 OXLP significantly reduced their expression (Figure 3A and 3B). The intestinal epithelium contains tight junctions that link neighboring cells to create a barrier preventing the free flow of substances between cells [36]. Tight junctions are made up of proteins such as occludins (OCLN) and claudins (CLND). Results showed that stimulation of the intestinal epithelium with LPS or OGD decreased the expression of *OCLN* and *CLND*, whereas co-treatment with HM sEVs or ω3 OXLP increased their expression (Figure 3A and 3B). We validated this by immunofluorescence. LPS and OGD treatment decreased the expression of tight junction proteins (E-cadherin (E-CADH) in red and occludin in green), whereas co-treatment with HM sEVs or ω3 OXLP restored their expression and the architecture and cohesion of the intestinal epithelium (Figure 3C and 3D).

**Fig. 3.**
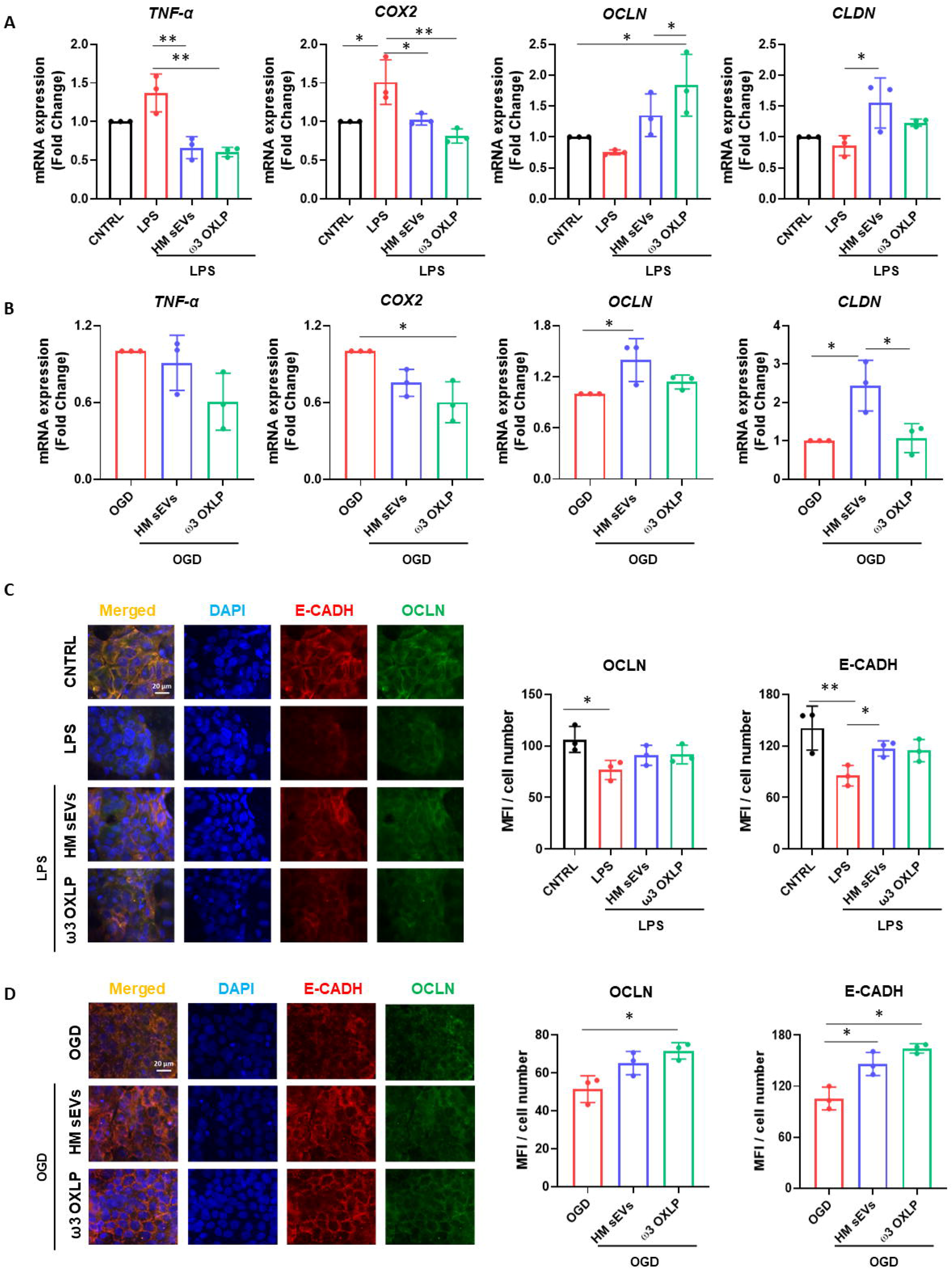
HM sEVs and ω3 OXLP dampen inflammatory responses in the inflamed epithelium. (A) Expression levels of *TNF-α*, *COX-2, OCLN* and *CLND* quantified by RT-qPCR in intestinal cells stimulated with lipopolysaccharides (LPS) and/or treated with 7.5 µg/mL sEVs or 0.5 nM of each of the three ω3 OXLP. (B) E-cadherin (E-CADH, red) and occludin (OCLN, green) immunofluorescence and nuclei staining (blue) show the distribution of tight junctions in the cell membrane. Unstimulated intestinal cells were used as controls. (C) Expression levels of *TNF-α*, *COX-2, OCLN* and *CLND* quantified by RT-qPCR in intestinal cell cultures under oxygen/glucose deprivation (OGD) condition and/or treated with 7.5 µg/mL sEVs or 0.5 nM of each of the three ω3 OXLP. The expression level of the target gene in each sample was normalized to *GAPDH* expression. (D) E-cadherin (E-CADH, red) and occludin (OCLN, green) immunofluorescence and nuclei staining (blue) show the distribution of tight junctions in the cell membrane. Scale bar: 20 µm. The bar graph shows the quantification of the mean fluorescence intensity (MFI). The graph represents the mean ± SD of three independent experiments. One-way ANOVA was used for statistical analysis. * p < 0.05, ** p < 0.01

### 3.5. Modulation of Pro-fibrotic Genes and Inhibition of Fibroblast Migration by HM sEVs and ω3 OXLP

Fibrosis is a pathological feature of most chronic inflammatory diseases, whereby fibroblast proliferation and migration lead to the excessive deposition of fibrous connective tissue, reducing its functionality [39]. LPS activates fibrosis, modulating the release of inflammatory cytokines and increasing fibroblast proliferation and migration [40], [41]. Results showed that the expression of the pro-inflammatory genes *TNF-α*, transforming growth factor beta (*TGF-β*), interleukin (*IL*)*-1* and *IL-6* increased significantly 24 h after LPS stimulation of fibroblasts. Treatment of LPS-activated fibroblasts with 7.5 µg/mL of HM sEVs or 0.5 nM of each of the three ω3 OXLP decreased the expression of these genes significantly (Figure 4A). In addition, the levels of other classical pro-fibrotic genes, toll-like receptor (*TLR*)*-4* and matrix metallopeptidase (*MMP*)*1,* were higher after LPS stimulation, and their expression was normalized after HM sEVs or ω3 OXLP treatment (Figure 4A). To test whether the changes in gene expression correlated with an anti-fibrotic response, the effect of HM sEVs and ω3 OXLP on fibroblast migration was assessed in scratch-wound assays. Stimulation with LPS promoted fibroblast migration and wound closure (37.8 ± 8.4%) of free area in LPS-treated cultures *vs* (57.5 ± 4.5%) in control cultures at 24 h. Contrastingly, the addition of HM sEVs and ω3 OXLP to LPS-activated fibroblasts reduced their migration, reaching levels similar to control cultures (Figure 4B).

**Fig. 4.**
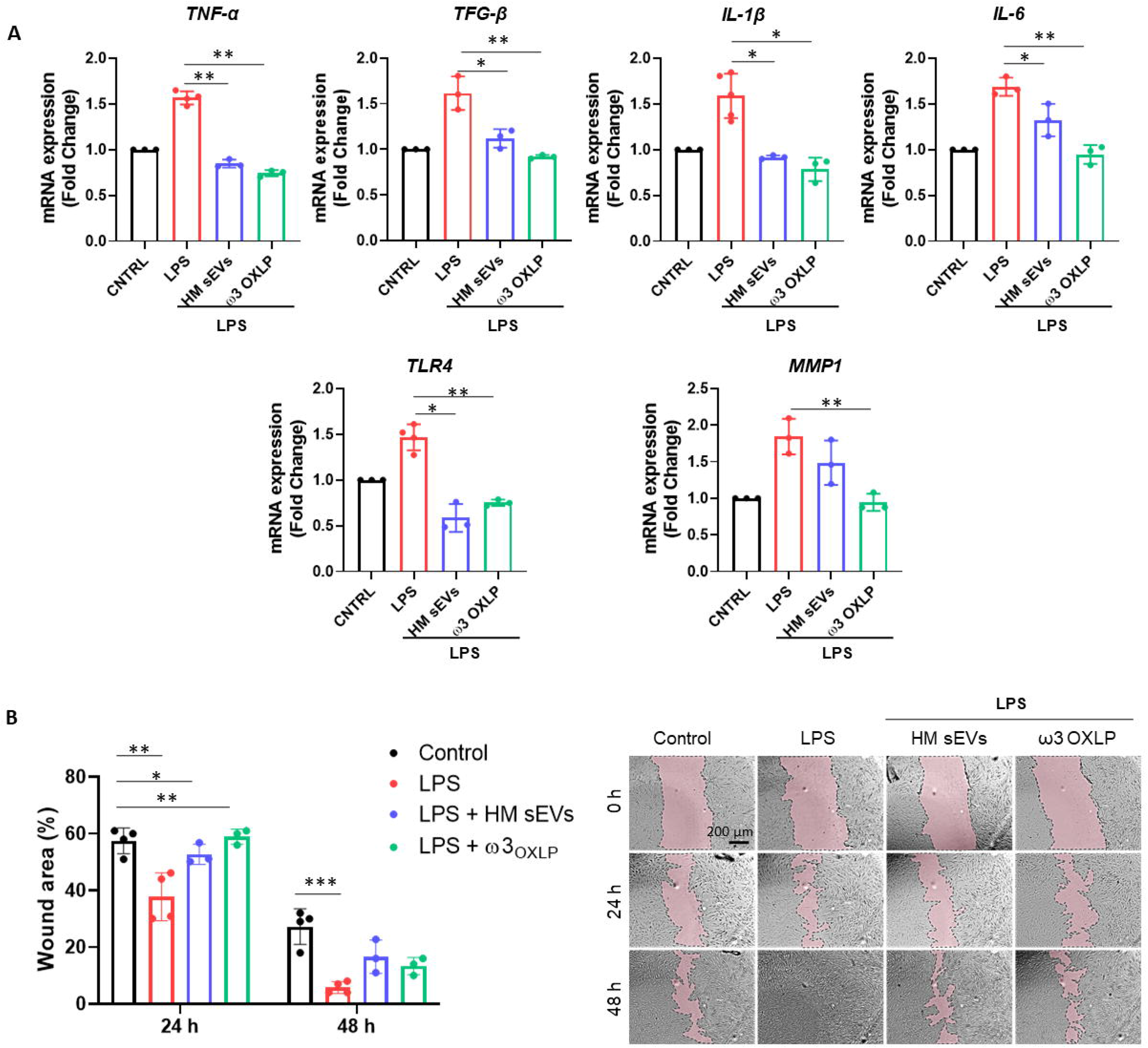
HM sEVs and ω3 OXLP prevent LPS-induced fibrosis. (A) Expression levels of *TNF-α*, *TGF-β*, *IL-1β*, *IL-6*, *TLR4* and *MMP1* quantified by RT-qPCR in fibroblasts stimulated with lipopolysaccharides (LPS) and/or treated with 7.5 µg/mL sEVs or 0.5 nM of each of the three ω3 OXLP. Unstimulated fibroblasts were used as controls. The expression level of the target gene in each sample was normalized to *GAPDH* expression. represented as mean percentage ± SD (B) Quantification of fibroblast wound closure at 24 and 48 h. Data were normalized to initial wound area and are represented as mean percentage ± SD. Representative brightfield images of wound healing assay at different times (0, 24 and 48 h) after wound generation on a monolayer fibroblast culture stimulated with LPS alone or treated with 7.5 µg/mL sEVs or 0.5 nM of each of the three ω3 OXLP. Images were taken at 10× magnification. Scale bar: 200 µm. Experiments were performed in triplicate. One-way ANOVA was used for statistical analysis. * p < 0.05, ** p < 0.01, *** p < 0.001.

### 3.6. Effects of HM sEVs and ω3 OXLP on Inflammatory Signaling Pathways, T-Cell activation, and Macrophage Polarization

Immune system cells, and more specifically macrophages, play a pivotal role in the pathogenesis of NEC, orchestrating both the inflammatory response and tissue repair processes [42], [43]. To study the effect of HM sEVs or ω3 OXLP on immune system cells, we performed different *in vitro* assays. First, 7.5 µg/mL of HM sEVs or 0.5 nM of each of the three ω3 OXLP were added to PBMCs to test whether they generated an immune response, activating the upregulation of pro-inflammatory cytokines genes *TNF-α*, *IL-1β*, *IL-6*, and *IL-8*. Results showed that ω3 OXLP did not activate proinflammatory signaling pathways with respect to non-stimulated control PMBCs, indicating that they are not immunogenic. However, the addition of HM sEVs resulted in a slight increase in the expression of *IL-1β*, *IL-6* and *IL-8* in PBMCs, although to a lesser extent than LPS (positive control) (Figure 5A). Second, we developed a T-cell activation and proliferation assay. Addition of ω3 OXLP to T-cells caused a slight reduction in their proliferation, whereas HM sEVs treatment appeared to increase proliferation (Figure 5B). Third, to study the ability of HM sEVs or ω3 OXLP to modulate Mϕ polarization, we differentiated monocytes to Mϕ type 1 (Mϕ1, pro-inflammatory) or type 2 (Mϕ2, pro-resolutive). During the differentiation to Mϕ1, some cultures were treated with HM sEVs or ω3 OXLP and surface markers were compared against non-treated Mϕ1 and Mϕ2 by flow cytometry. Results showed that the percentage of CD14^+^CD163^+^ cells, representative of a classical Mϕ2 phenotype, was not modified by HM sEVs or ω3 OXLP treatment (Figure 5C). Contrastingly, when the expression of cell surface receptors on differentiated and LPS-stimulated Mϕ1 were analyzed, we observed that treatment with HM sEVs significantly reduced the expression of the co-stimulatory molecules CD80 and CD86, and also HLA-DR expression to levels seen in Mϕ2. Treatment with the ω3 OXLP also reduced the expression of all three markers, although to a lesser extent (Figure 5C). To confirm the ability of HM sEVs and ω3 OXLP to induce Mϕ polarization, we measured the levels of proinflammatory TNF-α and anti-inflammatory IL-10 cytokines in the culture medium of Mϕ. Mϕ1 released a large amount of TNF-α and low levels of IL-10, and the opposite occurred with Mϕ2 (Figure 5D). Treatment of Mϕ1 with HM sEVs resulted in a profile more similar to Mϕ2, with a reduced amount of TNF-α and a higher amount of IL-10; and treatment with ω3 OXLP significantly reduced released TNF-α but failed to alter IL-10 release by Mϕ1 (Figure 5D).

**Fig. 5.**
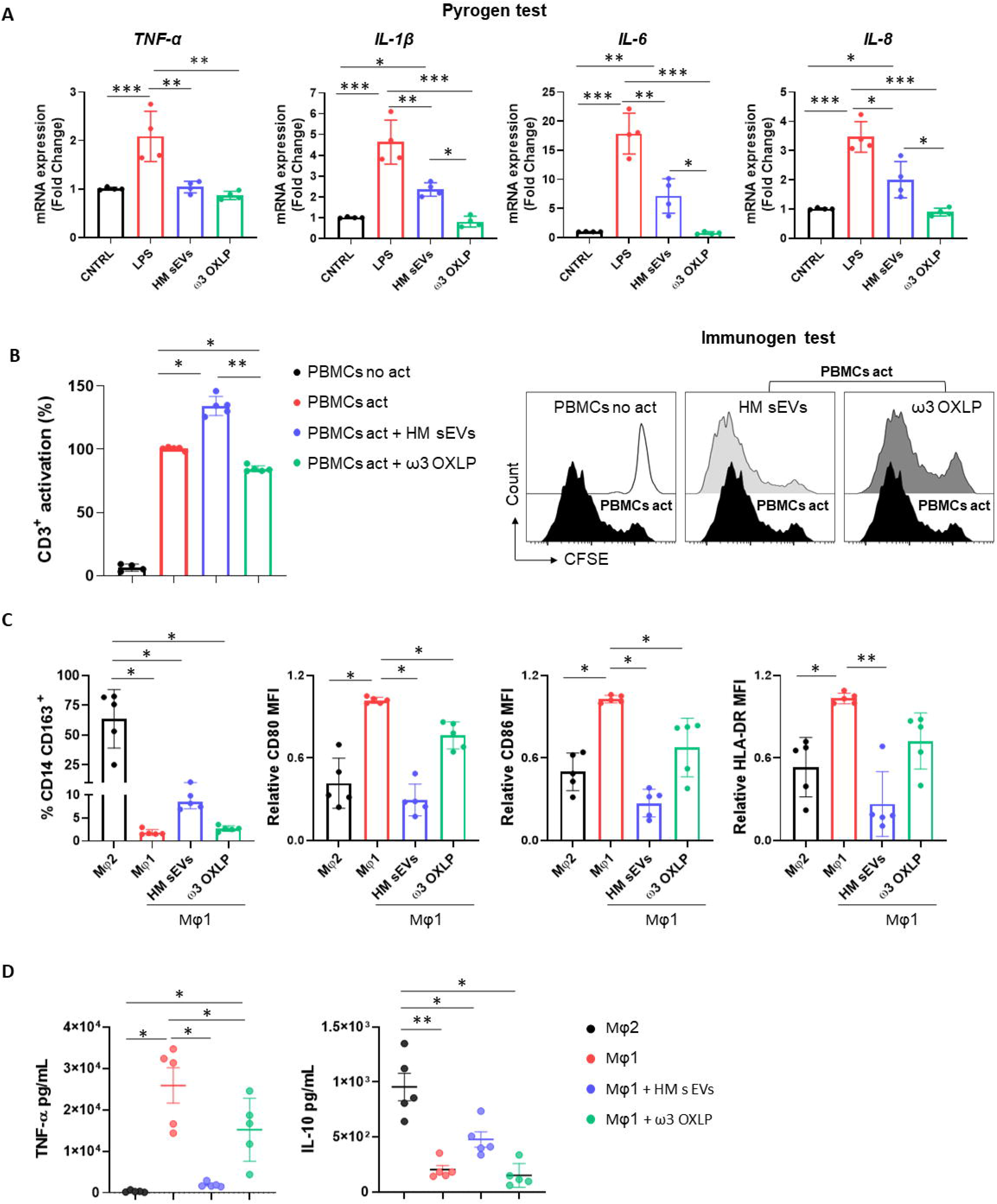
Response of HM sEVs and ω3 OXLP on immune system cells. (A) Expression of proinflammatory genes (*TNF-α*, *IL-1β*, *IL-6* and *IL-8*) in peripheral blood mononuclear cells (PBMCs) cultured for 6 h with treatments (HM sEVs and ω3 OXLP). Unstimulated and lipopolysaccharides (LPS)-stimulated PBMCs were used as negative and positive controls, respectively. The expression level of the target gene in each sample was normalized to *GAPDH* expression. The graphs represent the mean ± SD of four independent experiments. (B) PBMCs were stained with carboxyfluorescein succinimidyl ester (CFSE) and stimulated with anti-CD3 and anti-CD28 in the presence or absence of HM sEVs or ω3 OXLP. After 5 days, cells were stained with anti-CD3 antibody and T-cell proliferation was determined by flow cytometry measuring CFSE dilution. Suppression (percentage) was calculated from the expansion index. The graphs represent the mean ± SD of four independent experiments. Representative histograms are shown. (C) Monocytes were differentiated to Mϕ1 with treatment (HM sEVs and ω3 OXLP). Differentiation to Mϕ1 and Mϕ2 was used as a reference of pro-inflammatory and pro-resolving macrophages, respectively. After 5 days of differentiation, the percentage of CD14^+^ and CD163^+^ cells was assessed by flow cytometry. After LPS activation, CD86, CD80 and HLA-DR expression was assessed by flow cytometry. The mean relative fluorescence intensity (MFI) was calculated by dividing all individual data by the mean expression in Mϕ1. (D) TNF-α and IL-10 production by Mϕ was determined by ELISA 16 h after LPS stimulation. Graphs represent the mean ± SD of five independent experiments. One-way ANOVA was used for statistical analysis. * p < 0.05, ** p < 0.01.

### 3.7. Therapeutic Potential of HM sEVs and ω3 OXLP in an Experimental Model of Inflammatory Bowel Disease

The evident beneficial effects of HM sEVs and ω3 OXLP *in vitro* motivated us to test their therapeutic potential in an IBD model using TNBS administered intrarectally to induce severe colonic inflammation in mice [44]. Balb/c mice were divided into four groups: a healthy sham group, an untreated TNBS group, a treated TNBS group with 50 µg of HM sEVs and a treated TNBS group with a cumulative dose of 1.5 µg of ω3 OXLP. Treatments were dissolved in 100 µL of PBS and were orally administered by gavage just after induction of acute colitis by TNBS and at 24 and 48 h later. The sham group was treated with 100 µL of vehicle (PBS). On the fourth day, mice were sacrificed, and the regenerative and anti-inflammatory effects of the treatments were assessed.

We monitored weight loss of mice across the experiment (Figure 6A). The sham group showed no weight loss, whereas the TNBS group lost almost 20% of their weight. Mice treated with HM sEVs and ω3 OXLP also showed weight loss; however, this stabilized on the third day, reaching a maximum of 10% loss at sacrifice (Figure 6B). Colon length was shorter in the TNBS group than in the sham group, whereas TNBS-induced mice treated with HM sEVs and ω3 OXLP showed protection against colon shortening (Figure 6C).

**Fig. 6.**
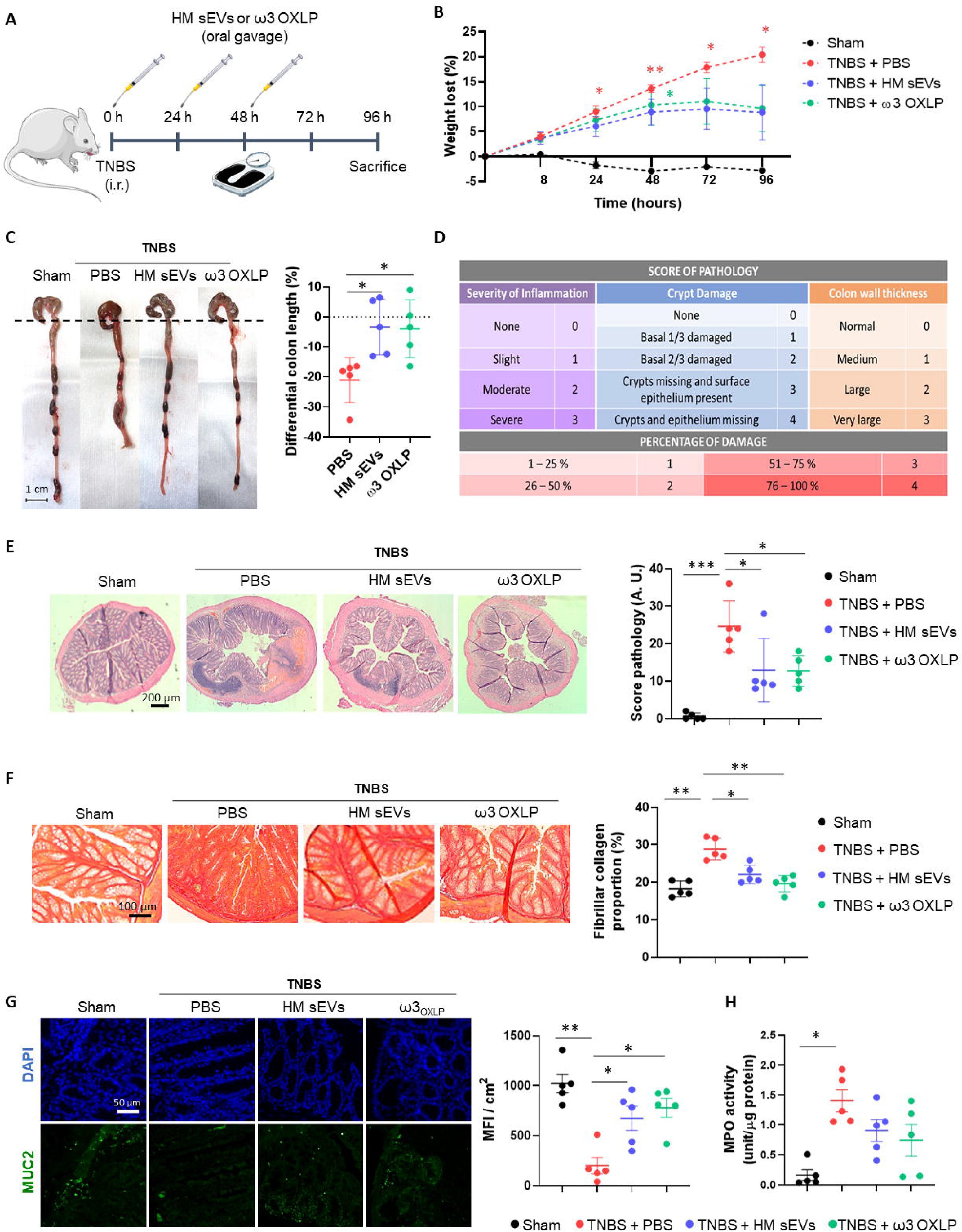
HM sEVs and ω3 OXLP attenuate disease in mice with TNBS-induced colitis. (A) Scheme of the *in vivo* experimental design. (B) Measurement of the weight loss of mice throughout the experiment (4 days). (C) Macroscopic images of colon tissue on day 4 after 2,4,6-trinitrobenzenesulfonic acid (TNBS) administration. Scale bar: 1 cm. Percentage differential length of the colon compared with the healthy group (horizontal dotted line). (D) Histology score table based on grade of pathology and percentage of damage. (E) Hematoxylin and eosin staining of representative histological sections of the colon of mice in the healthy group and in the PBS, sEVs and ω3 OXLP groups after TNBS administration. Scale bar: 200 μm. (F) Sirius Red staining was used to detect collagen fibers. Scale bar: 200 μm. Fibrillar collagen proportion (%) was calculated by dividing the area stained with red by the total tissue area. (G) Immunofluorescence of MUC2 (green) and nuclei staining (blue). Scale bar: 50 µm. Bar graph shows quantification of green mean fluorescence intensity (MFI) per cm^2^. (H) Myeloperoxidase (MPO) activity was measured in colon homogenates. Values were relativized by μg of protein tissue. The graph represents the mean ± SEM of five mice in each group. One-way ANOVA was used for statistical analysis. * p < 0.05, ** p < 0.01, *** p < 0.001.

Examination of colon histology revealed severe mucosal damage in the TNBS group, characterized by fewer intestinal glands, distortion of crypts and profuse inflammatory cell infiltration. By contrast, the TNBS group treated with HM sEVs and ω3 OXLP showed preserved tissue architecture with significantly less histopathological damage (Figure 6D and E). To investigate the mechanisms underlying colitis recovery after treatment with HM sEVs and ω3 OXLP, we used Sirius Red staining to assess tissue collagen fiber content. Intestinal fibrosis occurs in the context of inflammation, leading to tissue damage and impaired tissue reconstruction with marked deposition of extracellular matrix [45]. As expected, the percentage of collagen in the colon was significantly higher in the TNBS group than in the sham group, whereas the groups treated with HM sEVs or ω3 OXLP showed significantly lower levels of collagen (Figure 6F), indicating that treatment with HM sEVs and ω3 OXLP alleviated intestinal fibrosis in colitis.

The intestinal mucosa is protected by a variety of glycoproteins known as mucins (MUC), which play a role in the mucociliary transport system by trapping pathogens in a mucin gel layer [46]. To further explore the protective effects of ω3 OXLP from HM sEVs in experimental colitis, we investigated the expression of mucin-2 (MUC2) by immunofluorescence. Results demonstrated that treatment with HM sEVs or ω3 OXLP maintained MUC2 expression in TNBS-induced mice (Figure 6G). Because a correlation between disease severity in IBD patients and neutrophil infiltration has previously been reported [47], we used a MPO assay to assess neutrophil activity. MPO activity was significantly higher in the TNBS group than in the sham group, and treatment with HM sEVs or ω3 OXLP resulted in a trend for decreased neutrophil activity (Figure 6H).

### 3.8. Modulation of Immune Response and Cytokine Expression by HM sEVs and ω3 OXLP in TNBS-induced Colitis

An imbalance between proinflammatory and anti-inflammatory immune cells and cytokines is a key characteristic of IBD, which hinders the resolution of inflammation. To assess the modulation of immune responses by HM sEVs and ω3 OXLP, we examined cytokine expression in colon tissues of treated mice. Cytokine protein arrays revealed that the levels of several cytokines were higher in the untreated TNBS group than in the sham group, including intercellular adhesion molecule (ICAM)-1, tissue inhibitors of metalloproteinase (TIMP)-1, CC motif chemokine ligand (CCL)2, CXC motif chemokine ligand (CXCL)9, CXCL13, CXCL1, IL-1β, triggering receptor expressed on myeloid cells (TREM)-1, IL-1α, CXCL11, IL-17, and TNF-α. By contrast, the TNBS group treated with HM sEVs or ω3 OXLP showed significantly lower levels of these cytokines, with some approaching the levels seen in the sham group. Notably, the colitis-induced group treated with HM sEVs or ω3 OXLP had elevated levels of anti-inflammatory cytokines such as IL-10 and IL-1 receptor antagonist (IL-1Ra) (Figure 7A).

**Fig. 7.**
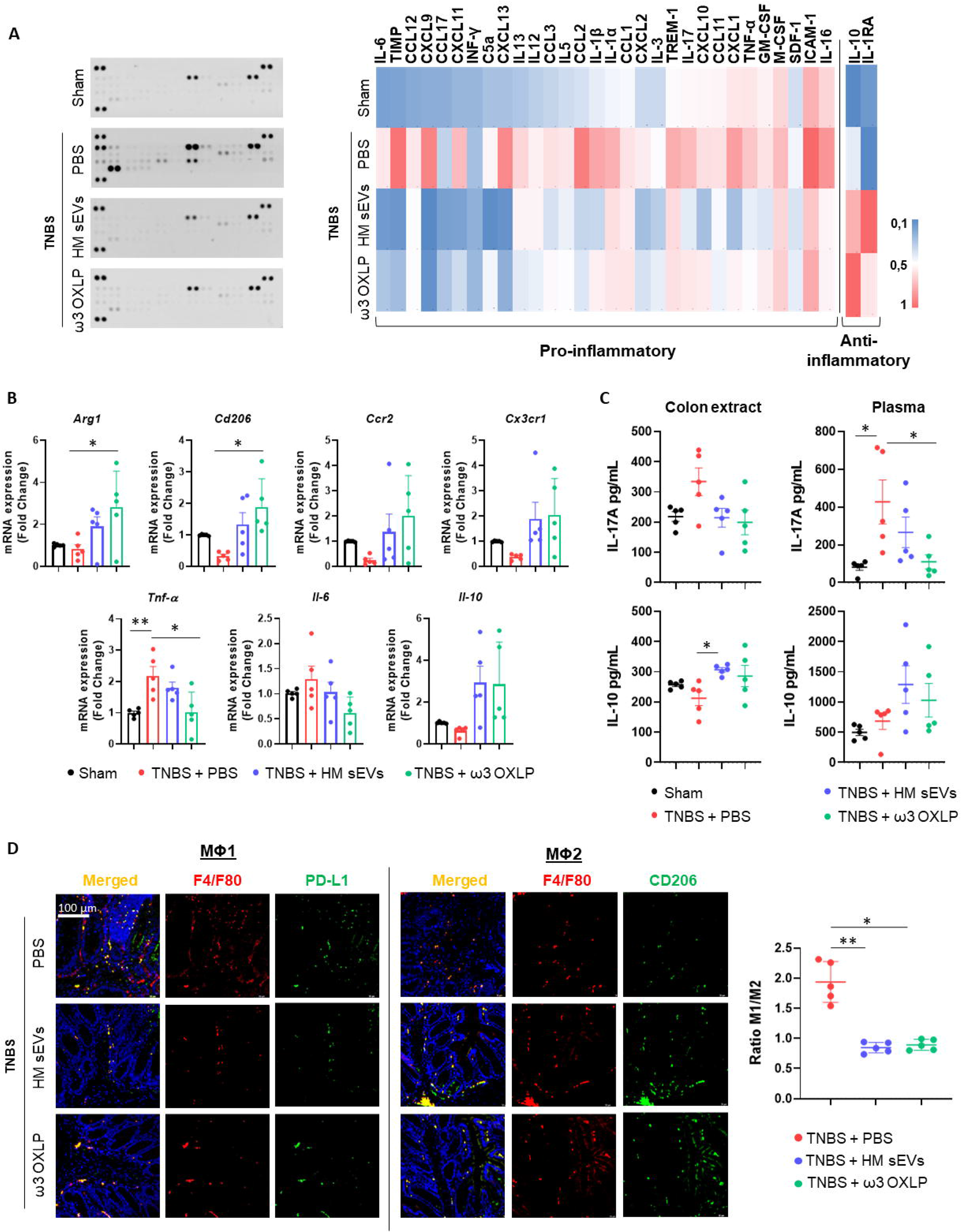
HM sEVs and ω3 OXLP change the ratio of infiltrating Mϕ1/Mϕ2. (A) Levels of inflammation-related cytokines were analyzed in colonic tissues by immunoblot array (left). Different time exposition was used to reveal different amounts of protein. The relative expression of each cytokine was quantified and represented in a heat map (right); data are representative of a pool of five animals per group. (B) *ArgI*, *Cd206*, *Ccr2*, *Cx3cr1*, *TNFα*, *Il-6*, and *Il-10* mRNA expression levels quantified by RT-qPCR in colon. Sham group was used as a control. Expression level of the target gene in each sample was normalized to *β-actin* expression. Graphs represent mean ± SEM of fold change of five independent experiments. (C) ELISA assay to assess IL-17A and IL-10 production (pg/mL) in colon extracts and plasma. (D) Immunodetection of F4/F80 (pan-macrophage marker, red) and PD-L1 (Mϕ1, green) or CD206 (Mϕ2, green) in colon samples 4 days after TNBS-induced colitis. Scale bar: 100 μm. Quantification of double-positive cells per mm^2^. Ten sections of 0.14 mm^2^ per mouse were analyzed. Graphs represent the Mϕ1/Mϕ2 ratio ± SEM of five mice. One-way ANOVA was used for statistical analysis. * p < 0.05, ** p < 0.01.

To further evaluate the immune response, we examined immune cell infiltrates in colon tissue. mRNA expression levels of pro-inflammatory cytokines (*Tnf-α* and *Il-6*) were significantly lower in the groups treated with HM sEVs and ω3 OXLP than in the untreated TNBS group, whereas the opposite pattern was seen for the anti-inflammatory cytokine *Il-10*. Analysis of Mϕ2-associated genes: *Arginase* (*Arg*1)*, Cd206, CC motif chemokine receptor (Ccr2*)*, and C-X3-C motif chemokine receptor* (*Cx3cr1*) [48], also revealed an increase in the groups treated with HM sEVs and ω3 OXLP (Figure 7B). We also measured IL-17A and IL-10 in plasma and colonic tissue by ELISA. The pro-inflammatory cytokine IL-17A was elevated in the TNBS group, but its levels were lower in mice co-treated with HM sEVs and ω3 OXLP. Conversely, IL-10 levels were lower in the TNBS group but were increased in TNBS mice co-treated with HM sEVs or ω3 OXLP, both in plasma and colon extracts (Figure 7C).

To gain further insight into the impact of the treatments on macrophage infiltration at the injury site during the disease, we performed an immunofluorescence assay using the classical macrophage marker F4/F80, combined with CD274 or CD206 to distinguish Mϕ1 and Mϕ2, respectively. The results demonstrated that the ratio of Mϕ1 to Mϕ2 was significantly higher in the untreated TNBS group than in the sham group, whereas treatment with HM sEVs and ω3 OXLP reversed this ratio, decreasing Mϕ1 and increasing Mϕ2 (Figure 7D). Overall, our findings indicate that HM sEVs and ω3 OXLP can mitigate the inflammatory response in TNBS-induced colitis by regulating immune cell infiltration and cytokine expression.

## 4. DISCUSSION

HM is the best food for newborns and PIs, as it provides them with all the necessary nutrients in the right measures. Indeed, the World Health Organization recommends mothers to breastfeed infants for the first six months of life to achieve optimal growth, development, and health [49], and HM is an essential member of the complex biological system between mother and infant [50]. In cases where breastfeeding is not possible or not chosen, infant formula may be a suitable alternative. However, while milk formula may provide adequate nutrition, it does not contain the immunological factors and other bioactive components present in HM, which provide additional protection against illness and promote optimal development. Recently, there has been renewed interest in bioactive lipids, as oxidized metabolites of PUFAs (oxylipins) have been detected in HM [51]. Several oxylipins, especially those derived from ω-3 fatty acids (ω-3-PUFAs), have been found to have anti-inflammatory properties and might be protective against chronic diseases and inflammatory conditions [52], [53].

Recent clinical studies have demonstrated that formula feeding might constitute a risk for NEC in PIs [54]. In this sense, supplementing HM is warranted. Here, we comprehensively investigated the presence of oxylipins derived from HM-sEVs and their therapeutic potential in the setting of intestinal inflammation. Our results support the idea of incorporating a combination of pro-resolving lipid mediators in milk formulations.

We show that HM-derived sEVs are loaded with 14-HDHA, 17-HDHA and 19,20-DiHDPA, that are pro-resolutive metabolites derived from the ω-3 fatty acid DHA and, in addition, both 14-HDHA and 17-HDHA are precursors of SMPs; specifically, maresins and D-series resolvins, respectively [55]. This may have a biological significance when considering HM-sEVs as therapeutic vehicles. Pizzinat *et al.* [56] previously reported the presence of lipid mediators in EVs derived from cardiomyocytes and mesenchymal stromal cells. However, they find a different oxylipin profile to the one found in HM-sEVs described in this work, probably because the source of EVs is different. Also, Chen *et al.* identified a total of 395 lipids in term and preterm HM-derived EVs [57], but no studies on oxylipins have thus far been reported.

We corroborated the utility of HM-sEVs for treating inflammatory disorders [58]. Since their discovery [59], the interest in the role of HM-derived EVs in early development has gained increasing interest, particularly with regards to their role in the gastrointestinal tract [14], and the contribution of HM-EVs to the maturation of the intestinal barrier has been studied in both physiological and pathological models [60], [61]. Moreover, recent studies have shown that HM EVs are resilient to digestion and can be endocytosed by intestinal epithelial cells [16]. In the present work, we show that HM-sEVs are taken-up by intestinal cells and that different damage stimuli (LPS or OGD) increase this process, pointing to a potential role for HM-sEVs in rescuing injured tissue from damage. In this regard, several studies have reported that milk derived EVs can ameliorate IBD in different *in vivo* models by suppressing immune cell infiltration and fibrosis, modulating MUC2 expression, reducing neutrophil activity, and promoting a pro-resolutive cytokine environment [62], [19], [58]. However, the potential use of milk-derived EVs is limited by the need for donors and the lack of scale-up procedures that would allow cost-effective commercialization.

We then tested whether ω3 OXLP present in HM-sEVs could reproduce the four main effects that are exerted by HM-EVs their selves in *in vitro* and *in vivo* models: (i) cell survival and proliferation, (ii) integrity (cell-cell junctions), (iii) resolution of inflammation and (iv) mucin production (additional defence) [14], [57], [18]. We demonstrate that ω3 OXLP present in HM-sEVs ameliorates oxidative stress and cytotoxicity in intestinal cells, resulting in improved cell viability and wound healing. Moreover, ω3 OXLP restored tissue integrity, increasing the expression of cell junction proteins including occludin, claudin and E-cadherin and halting fibrosis. ω3 OXLP was not immunogenic, endorsing its suitability for *in vivo* administration. Moreover, ω3 OXLP reduced T-lymphocyte proliferation and Mϕ1 polarization *in vitro*. It has been previously described that different SPMs can stimulate a switch in macrophage phenotype from a proinflammatory to a pro-resolving M2-like phenotype [63].

Several studies have addressed the potential beneficial effects of PUFAs in inflammatory diseases. For example, RvD1 administration (17-HDHA-derived) was found to reduce intestinal fibrosis in a colitis animal model [64]. In another study, Borsini *et al.* combined the ω3-PUFAs, EPA and DHA, to stimulate the production of lipid mediators, including 14-HDHA and 19,20-DiHDPA, which had neuroprotective effects. Also, treatment with ω3-PUFAs prevented neurogenesis loss and reduced apoptosis induced by pro-inflammatory cytokines in human hippocampal progenitor cells [65]. Regarding this latter strategy, increasing the intake of EPA and DHA provides the necessary substrates for the body to produce SPMs, which can be effective in boosting SPM levels indirectly and may have broader effects beyond the administration of specific SPMs. In this context, several studies have indicated that increasing the consumption of EPA and DHA can lead to higher concentrations of specific SPMs in human plasma or serum [66]. However, the relationship between the intake of EPA and DHA and the augmentation of particular SPMs remains unclear. The impact of EPA and DHA on SPM levels may be influenced by the minimum intake threshold of ω3-PUFAs required to stimulate significant endogenous biosynthesis of SPMs. While the availability of free EPA and DHA is crucial as substrates for endogenous SPM production, most of the EPA and DHA in the bloodstream, cell membranes, and intracellular compartments is esterified within complex lipids [67]. For this reason, the administration of ω3 OXLP rather than their precursors might overcome this problem. Interestingly, two of these OXLP are present in a commercial marine oil formulation, whose pro-resolutive properties have been demonstrated by our research group and others [33], [68].

The present study has several limitations that should be addressed. First, we did not use a NEC mouse model. NEC is the most common life-threatening gastrointestinal emergency experienced by PIs [69], affecting 7–8% of patients in the neonatal intensive care units and with mortality rates approaching 20–30% [70]. Nonetheless, despite the differences in their clinical presentation and affected demographics, emerging evidence suggests commonalities in the underlying inflammatory processes and molecular mechanisms between NEC and IBD, including dysregulated immune responses, mucosal barrier dysfunction, and altered gut microbiota composition, which contribute to intestinal inflammation in both NEC and IBD. While NEC primarily affects PIs, IBD encompasses a group of chronic inflammatory disorders that can occur in both children and adults. Moreover, the alterations in immune response, intestinal necrosis and fibrosis seen in the NEC model are relatively non-specific clinical manifestations that can be easily conflated with other gastrointestinal diseases, such as Crohn’s disease [71]. For this reason, we used the TNBS-induced mouse colitis model, as it shares common functional alterations with NEC [72].

A second major limitation is that although we detected other oxylipins, such as 9,10-DiHOME and 12,13-DiHOME (ω6-PUFAs), we did not analyze their therapeutic potential in our preclinical models. Nonetheless, the role of γ-linolenic acid (GLA), another ω6-fatty acid, was investigated recently in an elegant study on cardiac physiology [63]. The findings of these authors support the significance of ω-6 fatty acids in maternal milk, highlighting the complex interplay between specific fatty acids, such as GLA, retinoid x receptors, and the metabolic switch towards fatty acid utilization for energy production in cardiac myocytes after birth [73]. Further research exploring other ω6-PUFA-derived oxylipins in HM-sEVs and their therapeutic role in intestinal inflammation could provide valuable insights into the usefulness of these molecules in the resolution of inflammation.

Finally, although differential ultracentrifugation has been considering the gold standard for sEVs isolation, critical drawbacks of this technique include vesicle aggregation (especially originating from highly viscous solutions such as milk) and lipoprotein contamination, where high density lipids (HDLs) could sediment alongside HM-sEVs due to similar densities [74]. If the suspension has a large negative ζ potential, vesicles will tend to repel each other and there will be no tendency for be added [75].

In conclusion, oral administration of ω3 OXLP attenuates intestinal inflammation *via* inhibiting pro-inflammatory signaling pathways, restoring M2/M1 macrophage balance and preventing collagen deposition, preserving tissue integrity. Our findings support that a diet formula supplemented with this cocktail of ω3 OXLP may have great potential in protecting and preserving the gut health of PIs and adults with IBD.

## ACKNOWLEDGEMENTS

We thank Dr. M. Vento (Group of Perinatology at IIS La Fe) for providing human milk from healthy donors and Dr. Pilar Solves (Service of Hematology of *Hospital Universitari i Politécnic La Fe*) for providing buffy coats. We thank Rosa Vives for help in the *in vivo* experiments and histological procedures and Dr. Lusine Hakobyan for DLS analysis. We thank Kenneth McCreath for manuscript editing. The data presented herein were obtained using core facilities at IIS La Fe (Analytical Unit, Cytometry Unit, Cell Culture Unit, Animal Housing Facility and Microscopy Unit) and the Electron Microscopy Service at *Centro de Investigación Principe Felipe*. Pictures were drawn using images from Servier Medical Art., provided by Servier and licensed under a Creative Commons Attribution 3.0 unported license.

## STATEMENT

### Institutional Review Board Statement

The study was conducted according to the guidelines of the Declaration of Helsinki and approved by the Institutional Ethics Committee of Hospital Universitari i Politècnic La Fe (Approval Number 2021-071-1 & 2022-748-1).

### Informed Consent Statement

Written informed consent was obtained from all breast milk donors involved in the study.

### Data Availability Statement

All data generated and/or analyzed during this study are included in this published article and its additional files. All the data can be shared upon request by email.

## AUTHOR CONTRIBUTIONS

P.S., I.-T. and J.-K were responsible for funding acquisition. A-I.-T. and J.-K. were responsible for design, performance and analysis/interpretation of the lipidomic experiments. M.G.-F. and E.A.-P. were responsible for design, performance and analysis/interpretation of the in vitro experiments. M.G.-F. and E.A.-P were responsible for design, performance and analysis/interpretation of the in vivo experiments. M.G.-F. and P.S. conceived the study and participated in its design and coordination and wrote the manuscript. All authors have read and agreed to the published version of the manuscript.

---

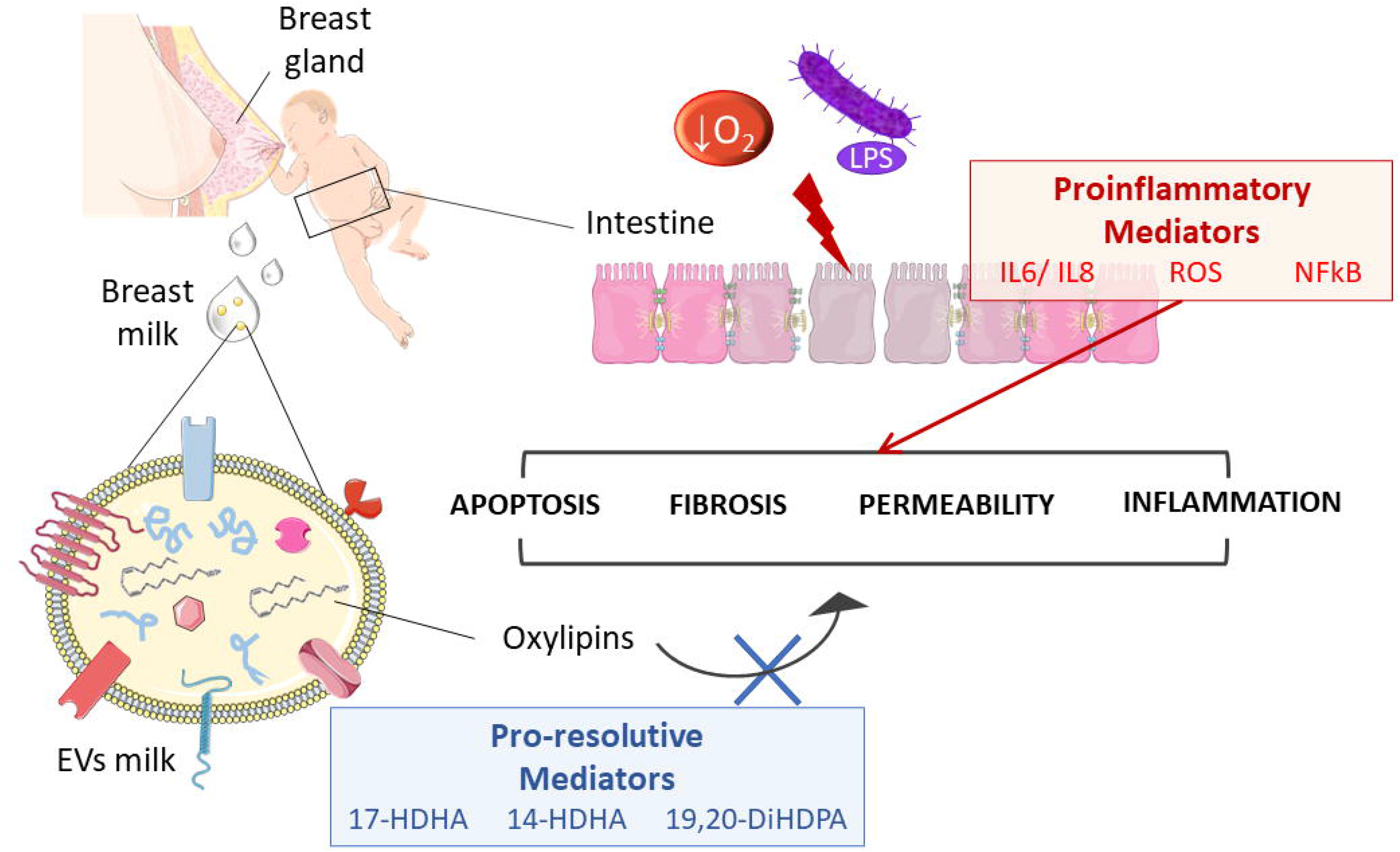

